# Quantitative imaging in whole-mount zebrafish embryos traces morphogen gradient maintenance and noise propagation in BMP signaling

**DOI:** 10.1101/2021.11.05.467413

**Authors:** Xu Wang, Linlin Li, Ye Bu, Yixuan Liu, Tzu-Ching Wu, David M. Umulis

## Abstract

Dorsoventral (DV) embryonic patterning relies on precisely controlled interpretation of morphogen signaling. In all vertebrates, DV axis specification is informed by gradients of Bone Morphogenetic Proteins (BMPs). We developed a 3D single-molecule mRNA quantification method in whole-mount zebrafish to quantify the inputs and outputs in this pathway. In combination with 3D computational modeling of zebrafish embryo development, data from this method revealed that Sizzled (Szl), shaped by BMP and Nodal signaling, kept a consistent inhibition level with Chordin (Chd) to maintain the BMP morphogen gradient. Intriguingly, BMP morphogen intrinsic expression is highly noisy at the ventral marginal layer in early zebrafish gastrula, where the gradient for DV patterning is established, which implies an unexpected role for noise in gradient shaping.

## Introduction

During embryonic development, morphogen gradients contribute to positional information for each cell and drive individual cell fate determination, meanwhile, stochastic noise can toggle gene expression levels (Hasty et al., 2000) and facilitate gene expression boundaries (Zhang et al., 2012). BMP ligands signal by binding to BMP receptors that form a higher-order receptor complex that phosphorylates the Smad5 transcription factor that binds to a co-Smad bef. BMP acts as a morphogen in numerous developmental contexts, including the patterning of the embryonic dorsal/ventral axis, neural tube, and Drosophila wing disc (Tuazon et al. 2020)(Bier and De Robertis 2015)(Briscoe and Small 2015). Szl, a downstream product of pSmad-mediated BMP signaling, also acts to regulate BMP signaling. Szl competitively binds proteinases (Tolloid and Bmp1) that target Chordin, an inhibitor of BMP signaling (Tuazon et al. 2020).

Meanwhile, the anterior-posterior position information is established by Nodal morphogen gradient secreted from the margin, inducing endoderm and mesoderm formation. It is the ratio of the Nodal to BMP that leads to the differentiation of cells, not the total amount of mRNA (Fauny, Thisse, and Thisse 2009). The same principle applies to the ratio of Smad2 and Smad5, which can direct the downstream Nodal and BMP signaling. Smad2 and Smad5 can selectively inhibit each other, making a diverse environment in which cells respond (Soh, Pomreinke, and Müller 2020). However, the significance of the mutual antagonism between Nodal and BMP signaling remains unclear.

RNAscope, the higher-resolution smFISH method, allows the detection of lower gene expression levels (Gross-Thebing, Paksa, and Raz 2014). Individual mRNA spots can be classified into nascent mRNA, which is brighter and more prominent in the nucleus, and mature mRNA, most of which are in the cytoplasm and smaller. By the threshold of spot intensity or watershed method, mRNA segmentations are well-used to quantify single cells or tissues (Mueller et al. 2013)(Carine Stapel et al., 2016). However, there are still a few issues remaining: firstly, how to distinguish mRNA spots from background or nonspecific binding; secondly, how to make sure each mRNA spot represents only one individual mRNA since some mRNA might be close to each other, incredibly high-level expressed genes; thirdly, how to distinguish between nascent and mature mRNA; fourthly, in order to elucidate the spatial noise, 3D mRNA segmentation in whole-mount zebrafish embryo is requisite.

Here, we observed staggered expression between *szl* and *chd* in the marginal region of zebrafish gastrula embryos. Applying high-resolution mRNA detection method to the input of mathematical modeling, we showed that Szl maintained the BMP signaling with a production region shaped by Nodal signaling. We investigated the intrinsic noise by comparing the variability among nascent mRNA; we revealed that BMP morphogen with high intrinsic noise is in the margin, suggesting stochastic noise may be involved in DV patterning, achieved by varying the cells fate.

## Results and Discussion

### Spatial detection of smFISH and protein at the single-cell level in whole-mount zebrafish embryos

To determine the relationship between input for secretion to the distributions of BMP signaling and BMP signaling activity, we used RNAScope method to simultaneously detect multiple individual mRNAs at the cellular level in whole-mount embryos. *bmp2b* mRNA began to be expressed at the zygotic stage, showing an obvious gradient pattern higher expression level in the ventral, whereas *chd* mRNA was expressed in the dorsal at 5.7hpf (Fig. 1A-C). *bmp2b* mRNA was detected at 2.5 hpf, with no *bmp2b* mRNA probes binding or background present, as a negative control (Fig. S1H-I). For positive control, we injected *venus-bmp2b* fusion mRNA into zebrafish embryos at the one-cell stage, followed by *bmp2b* and *venus* mRNA staining at 3hpf, when few endogenous *bmp2b* mRNAs were expressed. Based on observations, *bmp2b* mRNA was well colocalized with *venus* mRNA (Fig. S1 A-F). *szl*, as the target of BMP signaling, was also expressed in the ventral region (Fig. 1F-H). Nascent mRNA were observed exclusively in the nucleus and typically appeared as two or four big spots reflecting transcription activity, whereas mature mRNA appeared both inside and outside of the nucleus with smaller sizes, as shown in the diagram of Fig. 1J. We observed mature and nascent mRNA in whole-mount zebrafish embryos at the first time(Fig. 1D, E, I). *chd* nascent mRNA was easy to discern (Fig.1E) compared with *bmp2b* and *szl* mRNA, the active loci of which were challenging to recognize, suggesting that nascent mRNAs are more evident during higher transcriptional activity. Furthermore, we used *tld* mRNA as a positive control to detect the nascent mRNA. *tld* nascent mRNA was apparent in the EVL layer. When injected with α-amanitin, an inhibitor of RNA polymerase II and III, embryos only showed few *tld* mature mRNAs (Fig. S2E-F) compared to embryos injected with DMSO at 4.5hpf (Fig. S2C-D). However, it was hard to recognize the nascent mRNA of *bmp2b* by 2D imaging (Fig. S3A-B), prompting us to analyze bmp2b mRNA at the 3D level. Images were acquired using an upright confocal microscope with a 20x high NA (1.0) objective, which shows similar resolution to images taken on widefield microscopes (Fig. S1J, K).

**Fig. 1.**
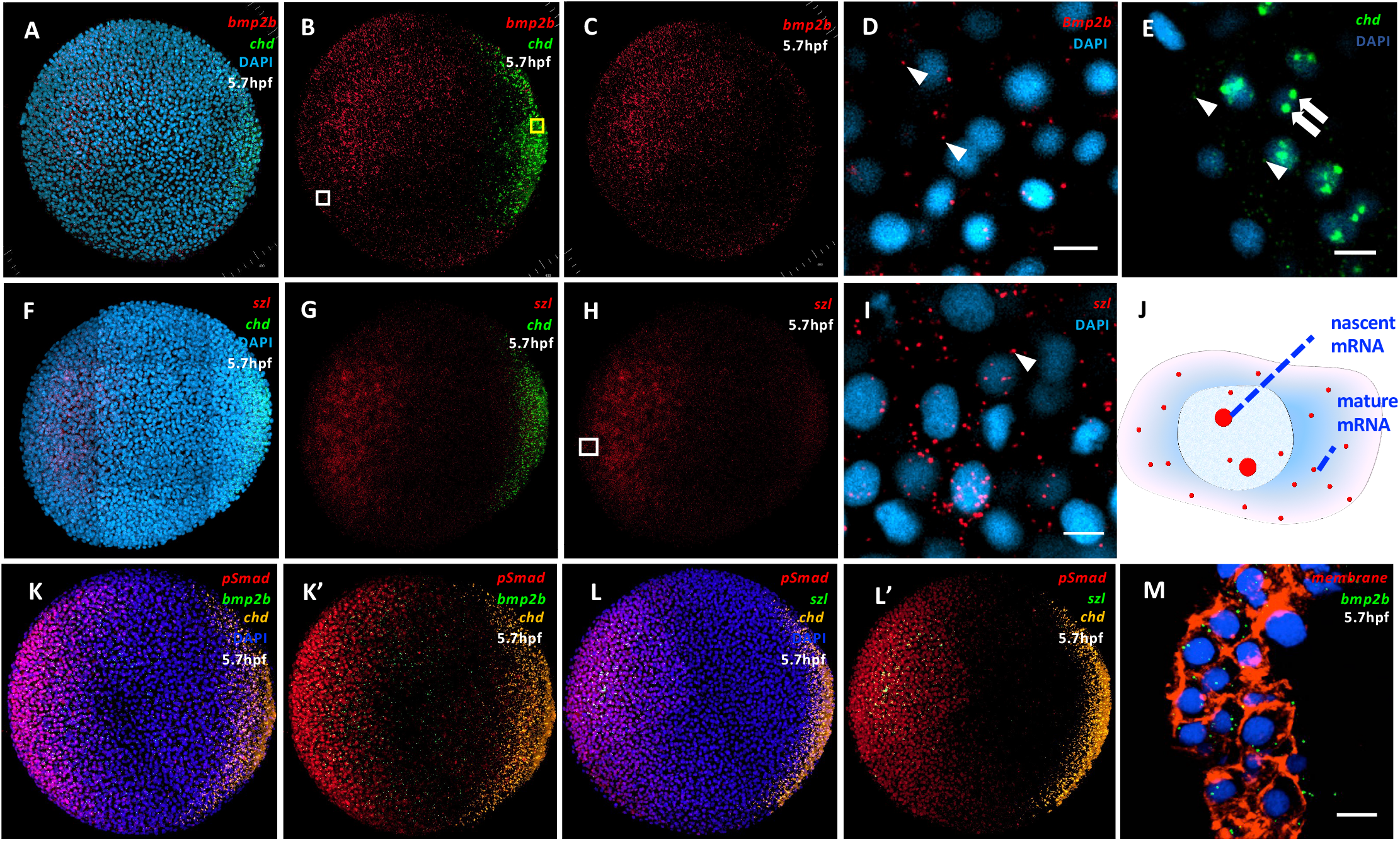
Specific detection of mature and nascent mRNA in zebrafish whole-mount embryos with RNAscope method. (A-E) *bmp2b* (red) and *chd* (green) single molecular mRNA expression imaged from animal view at 5.7hpf, with nucleus stained by DAPI (blue). (B) *bmp2b* and *chd* in A. (C) *bmp2b* in A. (D) White box in B at higher magnification. Arrowhead: *bmp2b* mature mRNA. Scale bar: 10 μm. (E) The yellow box in B at higher magnification. Arrowhead: *chd* mature mRNA; Arrow: *chd* nascent mRNA. Scale bar:10 μm. (F-I) *szl* (red) and *chd* (green) single molecular mRNA expression imaged from animal view at 5.7hpf, and nucleus are stained by DAPI (blue). (G) *szl* and *chd* in F. (H) *szl* in F. (I) White box in H at higher magnification. Arrowhead: *szl* mature mRNA. Scale bar: 10 μm. (J) example of nascent and mature mRNA. (K-L’) simultaneous expression of pSmad (red) and *bmp2b* (K, K’) or szl (L, L’) (green) single molecular mRNA at 5.7hpf, with nucleus stained by DAPI (blue). (M) membrane labeling (red) when detecting *bmp2b* mRNA (green) at 5.7hpf. Scale bar:10 μm.

Transcriptional responses to BMP signaling are mediated through pSmad. Comparison of BMP ligand levels with pSmad levels provides insight into positional information in this pathway. Comparison of pSmad levels with BMP target gene levels provides a measurement of transcriptional efficiency. Here, we demonstrate that we can detect pSmad and *bmp2b* (Fig.1K, K’) or szl (Fig. 1L, L’) mRNA in identical embryos without sacrificing the intensity of each. Cell membrane staining is critical for single-cell level detection of mRNA and protein because of its importance for image segmentation. Unfortunately, RNA scope protocol complicates this as F-actin structures of cell membranes are broken up during preparation. Remarkably we found that injecting membrane-labeling dye into zebrafish embryos at the one-cell stage circumvented this issue and provided membrane staining (Fig. 1M). Overall, we demonstrate the viability of staining single molecular mRNA and protein and cell membranes at the single-cell level; this method will enable the investigation of positional information and study the relationship between transcription factors and target genes, and noise at the cellular or embryonic level.

### Absolute quantification of nascent and mature mRNA transcripts in whole-mount zebrafish embryos

Previously mRNA quantification in zebrafish embryos was done by observing the mRNA quantity on 2D segmentation on cryo-sections (Carine Stapel et al. 2016). This approach loses the overall shape of individual mRNA of relatively large size. Some big spots like nascent mRNA and mature mRNA can show up on different slices of whole embryos during imaging, influencing each spot’s total intensity and volume matrix pixels. We developed 3D mRNA segmentation methods in whole-mount embryos following the flow chart shown in table 2, during which intensity drop-off correction is a prerequisite (Fig. S3 A-D). Transcription is mainly maintained in the interphase. Up to 4 transcriptional loci can show up in the nucleus during DNA replication (Fig.4F). This is consistent with similar findings in the mRNA segmentation of Drosophila embryos(Little, Tikhonov, and Gregor 2013) but was not observed in the 2D mRNA segmentation of zebrafish(Carine Stapel et al. 2016). To determine the individual mRNA candidates, we compared the *bmp2b* mRNA counts in entire embryos at 5.7hpf by image quantification to those from digital PCR (table 1). Representative transcriptional loci are shown in Fig 2B, with two big nascent mRNA spots inside of the nucleus. In digital PCR, the *bmp2b* mRNA probe targeted the middle of the *bmp2b* coding sequence. mRNA intensity is higher in mRNA strings transcribed more than half and vice versa, so, on average, the intensity of an individual nascent mRNA can be estimated as half of the averaged intensity of mature mRNA to match with the digital PCR results. Regardless of size and intensity, mature mRNAs were counted as one individual mRNA except for connected mature mRNAs, separated by local maximum (Fig 2C). The only variable, the intensity threshold, can be determined through fitting a curve from individual mRNA numbers by different thresholds (Fig. 2D), coupled with a digital PCR number (table 1, n=273778). We found that the ratio of maximum to averaged value, including volume matrix pixels (40/20) and intensity (10/5) of individual mRNA, was around 2 (Fig. S3 G, H), which is also found in previous work(Raj et al. 2008). Many individual mRNAs with a smaller area and lower intensity were also considered the actual single mRNA molecule shown in the peak of Fig. S3 G, H. Combining mRNA segmentation and digital PCR in whole embryos, background signal was authentically removed from smFISH images. Nascent and mature mRNA was divided based on location, size, and intensity.

**Table 1.**
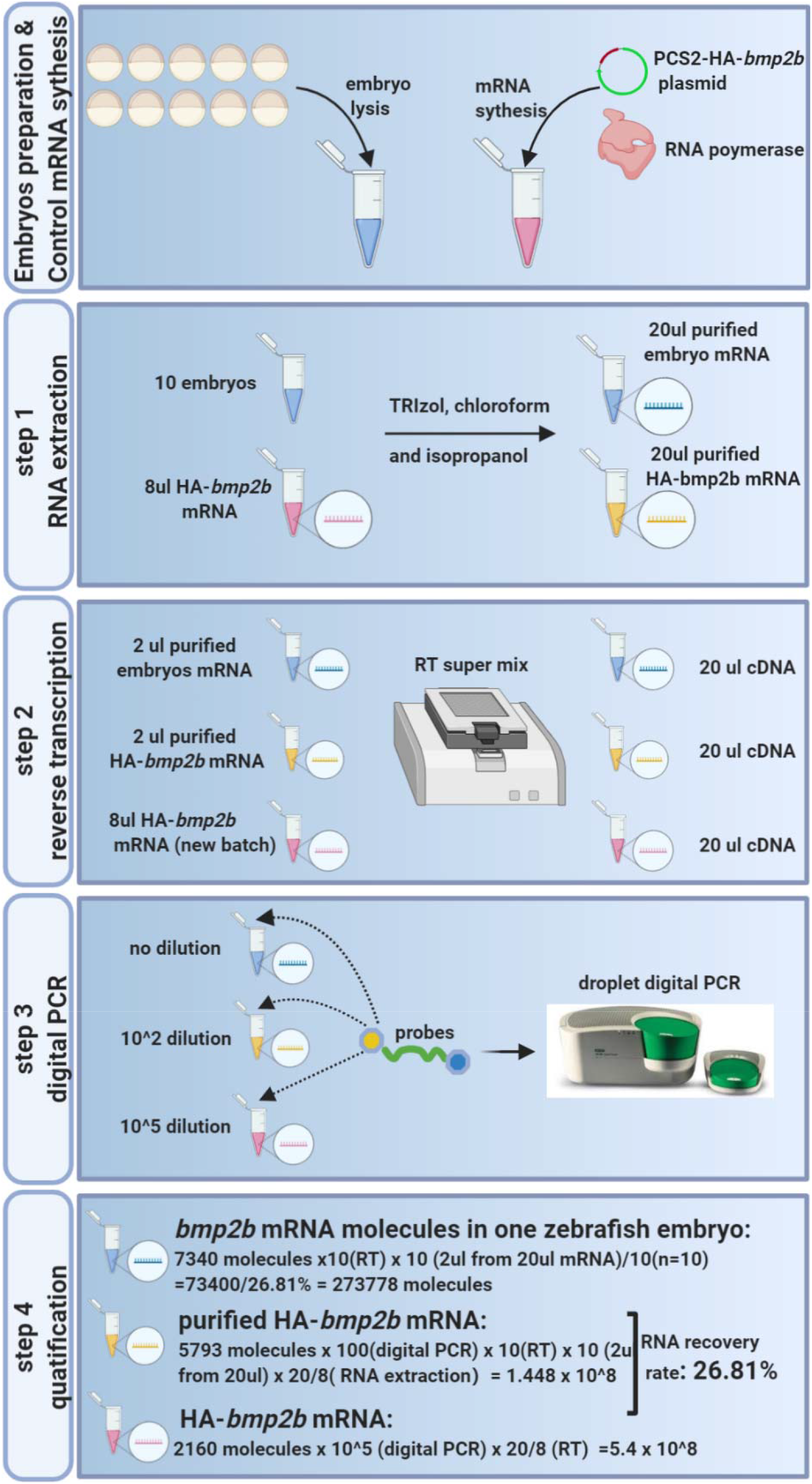
Quantifiy *bmp2b* mRNA number in one embryo by digital PCR method

**Table 2.**
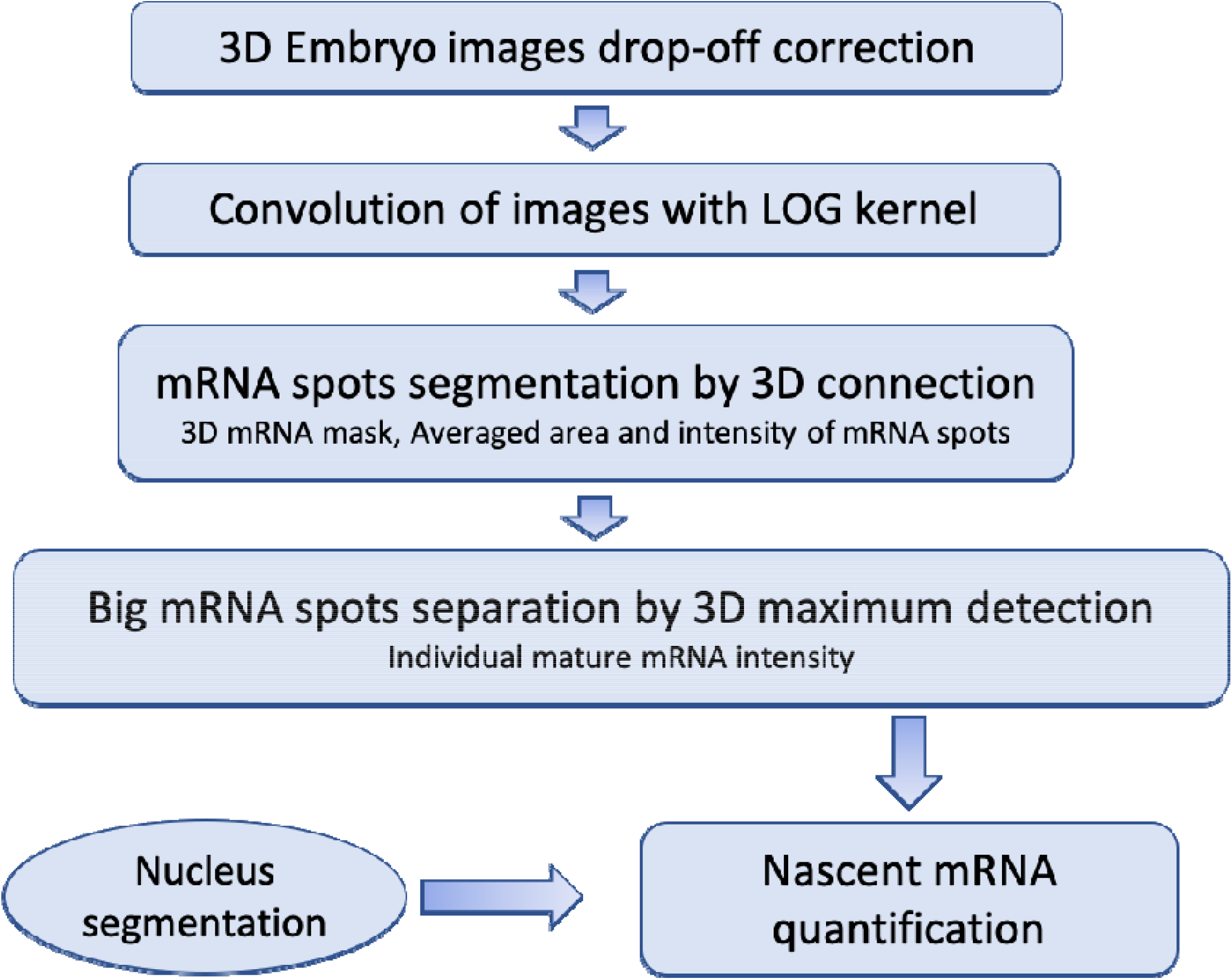
Workflow of mRNA_WME_quant (WME: whole mount embryo)

**Table 3.**
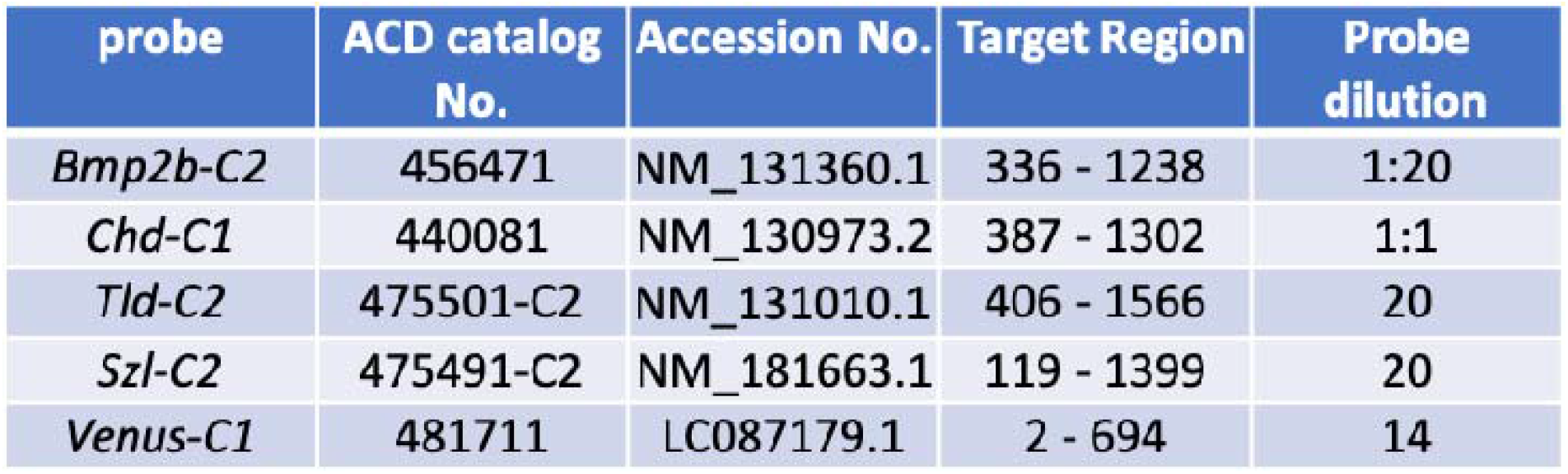
Information of probes in RNAscope method

| probe | ACD catalog No. | Accession No. | Target Region | Probe dilution |
| --- | --- | --- | --- | --- |
| *Bmp2b*-C2 | 456471 | NM_131360.1 | 336 - 1238 | 1:20 |
| *Chd*-C1 | 440081 | NM_130973.2 | 387 - 1302 | 1:1 |
| *Tld*-C2 | 475501-C2 | NM_131010.1 | 406 - 1566 | 20 |
| *Szl*-C2 | 475491-C2 | NM_181663.1 | 119 - 1399 | 20 |
| *Venus*-C1 | 481711 | LC087179.1 | 2 - 694 | 14 |

**Table 4.**
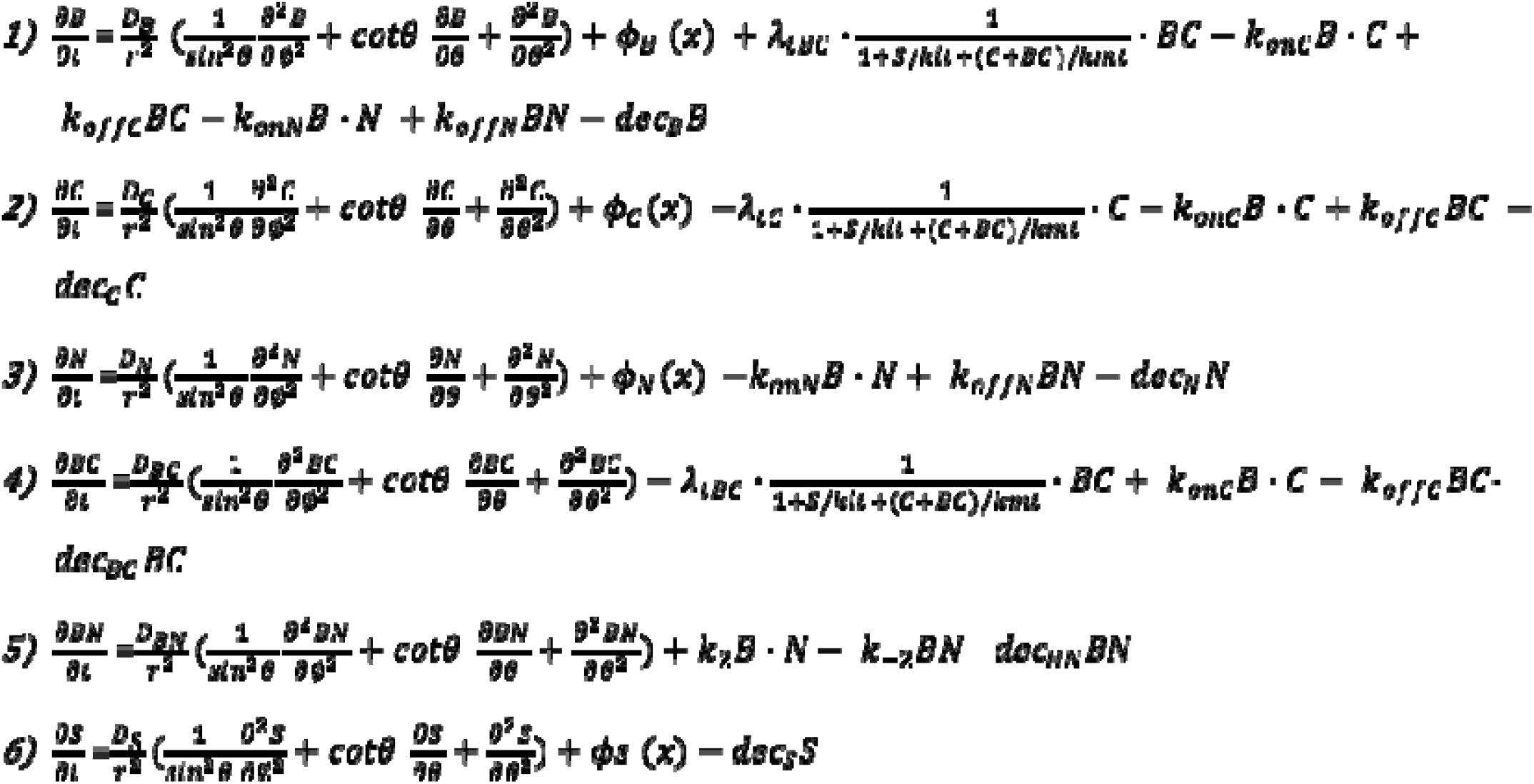
Mathematical model

**Fig. 2.**
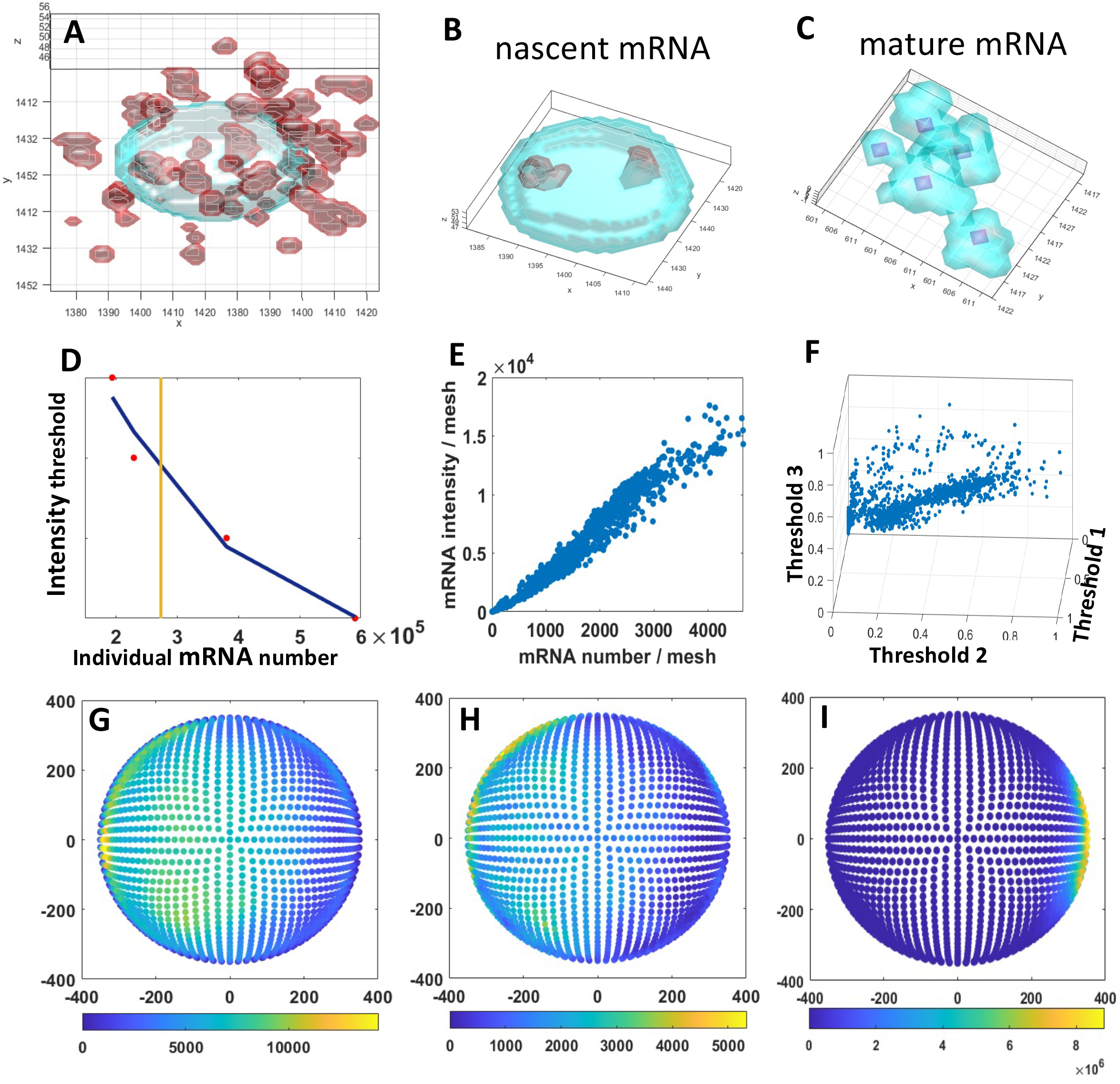
Quantification of *bmp2b* nascent and mature mRNA in whole mount embryo at 5.7hpf. (A-C) mRNA and nuclear segmentation with nuclear mask (light blue) and smFISH masks(red). (B)Nascent mRNA from (A). (C)Separation of connected cytoplasmic mature mRNA from (A). Purple dots indicate the local maximum of single cytoplasmic mRNA. (D) Individual mRNA number after entire image processing of single embryo with different intensity threshold. Scatter red dots represent the test value with blue linear fitting. The yellow line shows the individual mRNA number from digital PCR. (E)mRNA number and intensity scatter plot in whole-mount embryo. (F)mRNA number scatter plot at three intensity thresholds in whole-mount embryo. (G)*bmp2b* mRNA number distribution. (H) *bmp2b* mRNA intensity distribution. (I) *chd* mRNA intensity distribution.

### Determining critical factors in quantifying mRNA distribution across whole embryos

In order to detect the relationship between mRNA number and intensity at different positions on whole embryos, we established a whole sphere mesh to represent the zebrafish embryos at the blastula stage and mapped the mRNA spots to the corresponding mesh. Interestingly, total mRNA number and intensity were linear with each other at each mesh (Fig. 2E). The *bmp2b* mRNA 3D distributions across whole embryos are also similar between different mRNA numbers (Fig. 2G) and mRNA intensities (Fig. 2H) in each mesh, being more highly expressed in the ventral animal and marginal regions. Embryos were rotated into standard alignment using the *chd* mRNA expression (Fig. 2I) as a marker of the dorsal region. Notably, the mRNA number in each mesh is sensitive to the intensity threshold in mRNA segmentation (Fig. 2F). All of these results suggest that the most accurate and efficient way to visualize the smFISH distribution is to first determine the intensity threshold by combining 3D mRNA segmentation and digital PCR, then apply this threshold to 2D mRNA segmentation of more samples, which is around five times faster than 3D mRNA segmentation. Using the intensity from 2D mRNA segmentation, we effectively quantified averaged *bmp2b* (Fig. S3 E, F) and *szl* mRNA (Fig. 3E) distributions in more embryos. *bmp2b* mRNA production evolved from occurring highly in the ventral animal region at 4.7hpf (Fig. S3 E) to occurring highly in both the ventral animal and ventral marginal regions at 5.7hpf (Fig. S3 F), suggesting *bmp2b* starts to play a vital role in the margin between 4.7hpf and 5.7hpf.

**Fig. 3.**
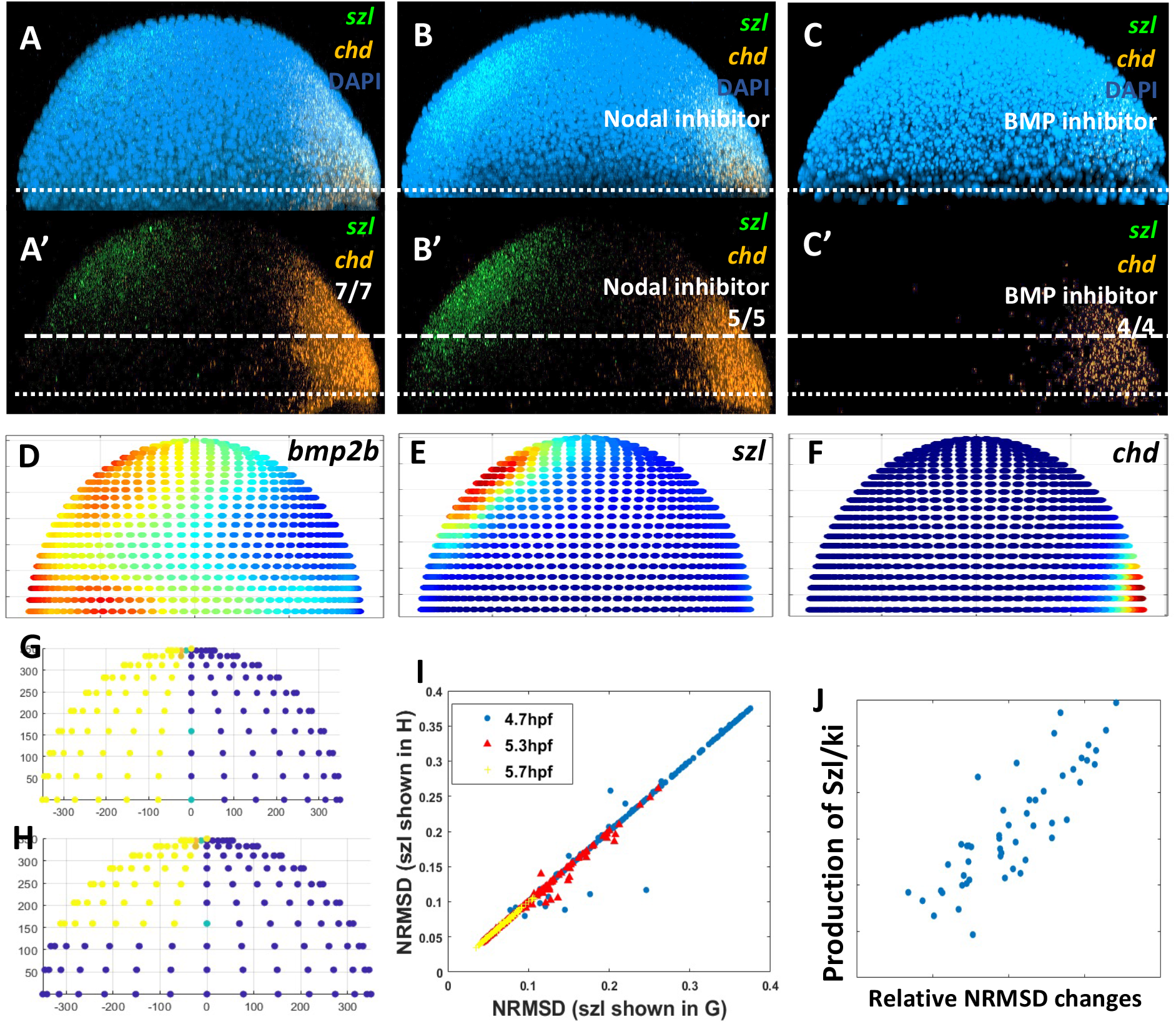
*Szl* mRNA expression maintaining the BMP morphogen gradient by BMP and Nodal signaling. (A-C’) *szl*(green) and *chd* (orange) single molecular mRNA expression imaged from lateral view at 5.7hpf, with nucleus stained by DAPI (blue). (A, A’) WT control. (B, B’) Nodal inhibitor. (C, C’) BMP inhibitor. (D-F) Averaged mRNA intensity distribution from lateral view at 5.7hpf.(D)*bmp2b*, n=4.(E)*szl*, n=7 (F) *chd*, n=7. (G-H) *szl* input in 3D mathematical modeling. (I) Scatter plot of NRMSD from different *szl* input in G and H. (J) Scatter plot of RNMSD and the ratio of *szl* production to ki.

### Szl maintains the inhibition level in the marginal layer by BMP and Nodal signaling

Higher-precision methods will give us a better resolution to detect the mRNA spatial distribution appropriate for mathematical modeling input. We found that *bmp2b* showed a gradient from ventral to dorsal in the ventral marginal and animal regions (Fig. 3D, Fig.S3 E). *chd*, as the target of Nodal signaling(Bradham et al. 2010), was expressed in the margin at 5.7hpf (Fig. 3F) and disappeared in the margin at 8hpf (Fig. S4 C,C’), which is consistent with Nodal expression from the blastula to gastrula stage(Rogers et al. 2017). Interestingly, *szl*, as the target of BMP signaling, was only expressed on the animal excluded from the ventral margin (Fig. 3E) and not colocalized with pSmad in the margin when simultaneously detecting *szl* and pSmad in whole-mount embryos at 5.7hpf (Fig. S4 B, B’). Remarkably, *szl* recovered the expression in the margin at the gastrula stage (Fig. S4 C, C’). This finding is consistent with another BMP target gene, foxi1, induced by high levels of BMP and low Nodal levels (Soh, Pomreinke, and Müller 2020). We postulated that *szl* might be inhibited by Nodal, which is only expressed in the marginal layer at 5.7hpf. To test this, we utilized a Nodal inhibitor (SB-505124) and a BMP signaling inhibitor (LDN193189) to evaluate the role of Nodal and BMP signals on *szl* expression. Importantly, We found that application of nodal inhibitor lead to resumed expression of *szl* at the ventral margin (Fig. 3A, A’, B, B’), and *szl* was entirely blocked by BMP inhibitor (Fig. 3C, C’). These results suggest that BMP and Nodal cooperatively shape the expression of *szl*.

Previous work shows that BMP and Nodal signaling can selectively inhibit each other to specify the cell type(Soh, Pomreinke, and Müller 2020). Our data implied that this mutual inhibition might play an essential role in BMP morphogen maintenance. To determine the impact of different *szl* expression on a three-dimensional level, we modified our previously-built three-dimensional growing domain model, which includes the growth of epiboly and 3D patterning(Li et al. 2020); the mathematical equations for this model are shown in Table 4. We adopted 182 sets of parameters screened from our 10 million 1D simulations(Tuazon et al. 2020) and applied different expression regions of *szl* as simulation input, with margin (Fig. 3G) or excluded from the margin (Fig. 3H). We calculated the NRMSD of modeling output data and experimental pSmad data, representing the fitting error, on the margin with two different *szl* expression inputs. Strikingly, some of the NRMSD can be significantly reduced by taking out the *szl* expression in the margin (Fig. 3I) and closely related to the ratio of *szl* production rate to ki represents the strength of Szl protein suppression (Fig. 3J). These results suggest that *szl* and *chd* change the expression region in the margin where dorsal-ventral patterning is determined to maintain the inhibition level of BMP morphogen.

### BMP morphogen intrinsic noise implies the cell state

Understanding the stochastic nature of signaling response between different cells is a fundamental challenge in biology. Intrinsic noise mainly comes from elements involved in transcription rates and can be evaluated by two-factor assays (Elowitz et al. 2002) or the sizes of transcribing loci (Stapel, Zechner, and Vastenhouw 2017)(Little, Tikhonov, and Gregor 2013). Extrinsic noise originates from the differences among cells, such as cell cycle stage or differential abundance of transcriptional factors (Zopf et al., 2013). We can detect the intrinsic noise through the volume pixels and intensity of nascent mRNA by 3D mRNA segmentation. We observed nascent mRNA in 49% of all nuclei ranging from 1 to 4 loci (Fig.4F), implying that about half of the cells are in the interphase. Strikingly, we observed four nascent mRNA in the form of 2 pairs of sister chromatids, indicating DNA undergoing replication. Of the observed nuclei, 20% had only one nascent mRNA (Fig 4E, F), suggesting that the transcriptional activity is independent between two chromosomes. This could also happen when more than one nascent mRNA colocalizes with each other. In order to access the instantaneous transcriptional activity of the BMP morphogen, we evaluated the variation among the total intensity of each nascent mRNA in the nuclei. Importantly, we found that the intrinsic noise was much higher in the marginal region for nuclei with both two (Fig. 4A, B) and four nascent mRNAs (Fig. 4C, D). In a whole-mount embryo, the nascent mRNA expression region is similar to the *bmp2b* mRNA in a whole-mount embryo (Fig.3D), and only the marginal region is highly noisy, which facts were clearly shown in amplified figures (Fig.4B, 4D). Representative nascent mRNA of higher noise levels was shown in Figure 4E to be compared with the one with lower noise (Fig. 2B). Zebrafish embryos start to establish the DV patterning in the margin at 5.7hpf, where and when a consistent noisy pattern was observed for intrinsic BMP morphogen noise, which noise can represent the state of the cell across whole embryos.

**Fig. 4.**
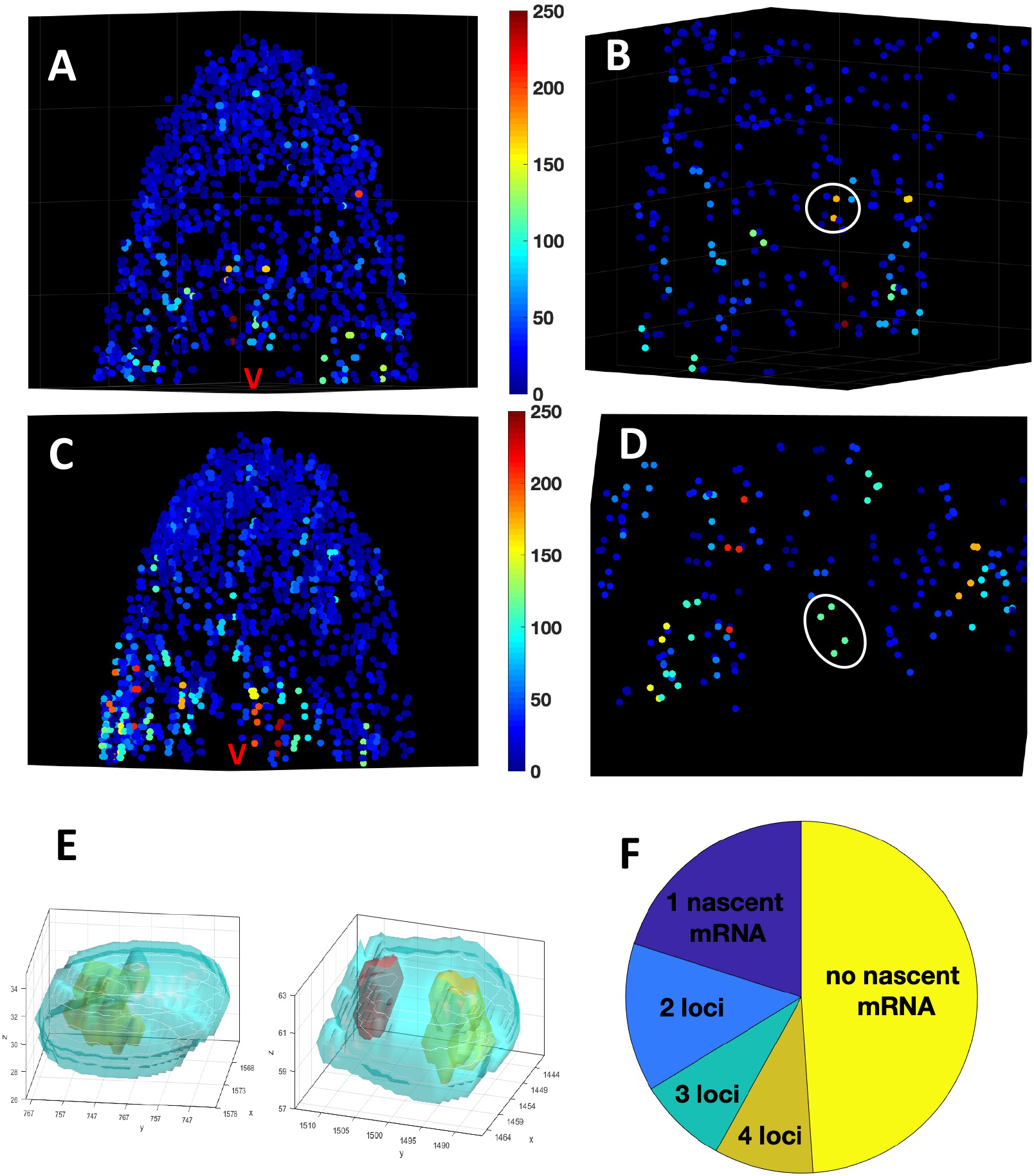
*bmp2b* Intrinsic noise indicating the cells state across the whole embryo at 5.7hpf. (A)Intrinsic noise distribution of nucleus with two nascent mRNA. The lateral view and ventral is in the middle. (B)Higher magnification from A. Representative two nascent mRNAs are marked with circles. (C)Intrinsic noise distribution of nucleus with four nascent mRNA. The lateral view and ventral is in the middle. (D)Higher magnification from C. Representative 4 nascent mRNAa in one nucleus are marked with circle. (E) Representative one nascent mRNA in the nucleus and two nascent mRNA in the nucleus with different sizes. (F)Statistic of nucleus number with different nascent mRNA number.

## Materials and methods

### Zebrafish

All procedures on zebrafish adults and embryos were approved by the Purdue University Institutional Animal Care and Use Committee (IACUC: 1501001180A004)). Wide-type (TL) embryos and Tg(HA: GFP) fish were collected in E3 water and fixed at the desired stages.

### smFISH using RNAscope method on cryosections

Zebrafish embryos are fixed with 4% PFA/PBS for 24 hours at 4°C. Wash the embryos with PBSTween, then dechorionated the embryos with forceps. Equilibrate the embryos into sucrose solutions: rinse first in 20% sucrose/PBS until all the embryos sink into the bottom, followed by 30% sucrose/PBS wash, wait until the embryos sink into the bottom. Then transfer the embryos into fresh 30% sucrose at 4°C. Embed embryos: Embed the embryos into Tissue-Tek® O.C.T. Use forceps to orient the embryo to ensure all the animal regions of the embryos facing the same direction. Blocks were quickly frozen in precooled isopentane on dry ice and stored at –80° C until performing cryosection. 10 µm sections were attached to *Superfrost* Plus microscope slides. Keep the slides at -80° C for at most two weeks. Samples were subjected to the protocol (Catalog Number 320293) from RNAscope based signal amplification (Advanced Cell Diagnostics). Samples were treated with Pretreat 2 and Pretreat 4 to enhance the accessibility of probes further. Mixed target probe hybridization was performed for 2 hours at 40°C. Four signal amplification systems then followed samples. DAPI was used at the last step to stain the nuclear and mounted with Fluoromount-G (SouthernBiotech, Cat NO 0100-01).

### smFISH using RNAscope method on whole-mount embryos

We used the protocol shown in our published paper(Li et al. 2020). Detailed probes information and dilution are shown in table 3. Make sure the embryos are in white, not yellow, color when air-dried from methanol.

### RNA staining using RNAscope followed by pSmad staining

We tested different staining methods: 1) mRNA staining first, then pSmad Immunostaining; 2) pSmad staining first, followed by mRNA staining. Test results showed that the first method was best. We made some modifications for the RNAscope staining part: RNA probe concentration was two times of original shown in table 1; RNA probes hybridization time is 12 hours. After the mRNA staining, embryos were penetrated in 0.1% Triton X-100 in PBS for 1 hour at RT. Embryos were blocked in blocking buffer (4% BSA(Millipore Sigma, #126615), 1% DMSO, 0.1% Triton X-100 in PBS) overnight at 4°C, and then stained with anti-phosphoSmad1/5/8 antibody (Cell Signaling Technology, #9511) at 1:100 diluted in blocking buffer overnight at 4°C. Then embryos were detected by goat anti-rabbit cross-adsorbed Alexa Fluor 647-conjugated antibody at 1:500 dilution with DAPI overnight at 4°C (Thermo Fisher Scientific, #A21244).

### Membrane labeling

Vybrant™ DiI Cell-Labeling Solution (Thermo Fisher, #V22885) was firstly injected into zebrafish embryos at one cell stage.

### Bmp2b-Venus mRNA injection

The Venus-Bmp2b fusion protein was synthesized by the same method as the previous paper (Zinski et al., 2017). 100pg *venus-bmp2b* mRNA was injected into one-cell stage embryos, and embryos were fixed at 3hpf and followed by the Venus probe staining.

### Transcription inhibition

Inject 2nl of 0.2mg/ml alpha-amanitin (Sigma, #A2263) into the yolk at 4.5hpf. At the same time, inject 2nl MQ water into the yolk of the control groups. Fix all the embryos at 6hpf according to the developmental stage of the control group.

### Chemical Inhibitions

BMP signaling inhibitor LDN193189(Sigma, #SML0559), Nodal signaling inhibitor SB-505124 (Sigmal, #S4696), and GSK inhibitor also known as Wnt signaling activator BIO(Sigma, #B1686)were dissolved in DMSO to 10mM as the stock and diluted in E3 medium at 50 µM, 50 µM, and five µM respectively.

### Image Acquisition

Embryos were mounted in 1% low melting agarose on 35mm glass-bottom microwell-dish (Matek, P35G-1.5-10-C). Whole-mounted embryos were imaged with a 20×/1.0 Plan-Apochromat water immersion lens (D = 0.17 M27 75 mm) of Zeiss 800. *chd* mRNA was imaged by excitation 555nm wavelength. *bmp2b, tld*, and *szl* mRNA were imaged by excitation 647nm wavelength. z interval is set to 3μm, which is smaller than the pinhole thickness. Cryosections were imaged with a 63x oil objective of Zeiss 800.

### Absolute Quantitative Digital PCR in whole embryos

Ten zebrafish embryos were collected at 6hpf, dechorionated by placing in 1mg/ml of pronase, immersed and ground in Trizol, and precipitated by isopropanol and chloroform. 8μl of HA-*bmp2b* mRNA synthesized from pCS2(+)-HA-bmp2b plasmid(Little and Mullins 2009) was purified under the same conditions as the embryos group. RT reactions were performed in raw or purified HA-*bmp2b* mRNA and purified embryos mRNA using iScript™ Reverse Transcription Supermix (Bio-rad, # 1708840), followed by digital PCR with TaqMan hydrolysis probes using the QX100TM Droplet Digital PCR System. Primers are against the *bmp2b* open reading frame across the intron region: forward primer (Tm: 60.5): 5’-CCA GCA GAG CAA ACA CGA TA-3’, reverse primer (Tm:59.9): 5’-CAT CTC CGA GAA CTT GGT CC-3’. Hydrolysis probe (Tm:64.8) is between the amplicon: 5’-CTC CGC TGC GGA GCT GCG CA-3’. Template cDNA was diluted to the dynamic range of QX100 (from 1 to 120,000 copies/20μl reaction) shown in step3 of table 1. mRNA numbers from raw and purified HA-*bmp2b* mRNA digital PCR results are used to calculate the loss of mRNA, and the RNA recovery rate is 26.81%. We can quantify bmp2b mRNA number in one embryo by the equation shown in step4 of table 1.

### Individual cytoplasmic mRNA and nascent mRNA segmentation process

Image processing can be fulfilled by five major steps shown in table 2:

1. Drop-off correction on whole-mount embryos
2. Images convolution and filtering
3. mRNA spots segmentation by 2D or 3D connection
4. Big mRNA spots separation by 3D maximum detection
5. Nascent mRNA identification

### Drop-off correction on whole-mount embryos

Embryos were imaged with an upright objective in the same orientation with the marginal region facing to the bottom, making signal intensity reduced from the animal pole to the margin of embryos as light is scattered when signal detection deepening into the lower layer of the embryo the embryos. We assumed nucleus intensity was consistent on average in Tg(HA-GFP) zebrafish. We collected and fixed zebrafish embryos at 8hpf, followed by Immunostaining using an Anti-GFP antibody (Thermo Fisher, # A-11122) at 1:500 dilution. The second antibody was the same channel in *bmp2b* mRNA staining. Nucleus intensity in each layer of z-stacks was collected by the wavelet nucleus segmentation method in our lab(Wu et al. 2020) and averaged to represent the intensity in the z-direction. We use a linear equation to correct reduced-intensity: I_corrected= I_origin-z_position*dropoff_scale, in which dropoff_scale is -0.18 for 647 channel in 400μm thickness whole-mount embryos (n=5).

### Images convolution and filtering

Intensity corrected images were convolved with a Laplacian of Gaussian filter (LOG) of size 15 with a standard deviation of 1.5. The only varied parameter was the threshold for the smallest intensity in whole processing, and the yielding number was compared with digital PCR data. The z-stack interval (3μm) is smaller than the pinhole thickness(15.9μm) on the objective of our confocal, making each mRNA spot imaged more than one time at the same XY position during consecutive slices. To obtain remarkable mRNA total number, 3D mRNA segmentation in whole-mount embryos is indispensable.

### mRNA spots segmentation by 2D or 3D connection

We implemented two methods to identify mRNA spots of whole-mount embryos.

A. 2D mRNA segmentation. Potential particles were identified as 2D connected masks in each slice using the Matlab function bwlabel. 2D matrix was transformed to 3D matrix with z position, which can be used to quantify the intensity of mRNA spots around five times faster than 3D mRNA segmentation. However, mRNA number will be over quantified in embryos.
B. 3D mRNA segmentation. 2D matrixes from each slice were first combined to 3D matrix with z position. We identified the connected mask at the 3D level using the Matlab function bwconncomp. 3D mask and number of all the mRNA spots were saved for the next step.

### Big mRNA spots separation by 3D maximum detection

Cytoplasmic individual mRNA can connect to one big spot. Potential big spots were firstly extracted if the spots intensity and pixels volume are larger than 1.5 times the average value. We find the 3D local maximums, including location and maximum intensity in the surrounding area of spots by Matlab function nonMaxSupr of Piotr’s toolbox, in which we set the radius as 2 pixels based on the averaged pixels volume.

### Nascent mRNA identification

Nucleus segmentation was performed by the wavelet method. Nascent mRNA candidates were first selected if the spots intensity and pixels volume are larger than 1.5 times the average value; meanwhile, nascent mRNAs located inside the nucleus were identified by the nucleus and mRNA masks. The maximum number of nascent mRNA in each nucleus is 4. Some cytoplasmic mRNA may still transit in the nucleus with a smaller size and intensity.

### mRNA distribution visualization

To visualize the overall mRNA distribution thought out the embryo. Individual embryos’ mRNA data were mapped to a standard shape of partial sphere depending on the specific development stages (4.7hpf as 40% epiboly, 5.7hpf as 50% epiboly). The embryo was then rotated to the DV axis based on the distribution of Chd mRNA (located at the dorsal most). An averaged distributed sample points clouds were created through the sphere; the accumulated mRNA intensity was collected on each sample point based on a defined circle area around this point. Multiple mRNA distribution from the same development staged were then averaged to obtain a general distribution map of the mRNA of individual species.

## Code availability

smFISH segmentation and visualization codes are available for academic use as Supplementary data.

## ACKNOWLEDGEMENTS

We thank the help of Aasakiran Madamanchi for paper proofreading.

---

**Sup Fig.1.**
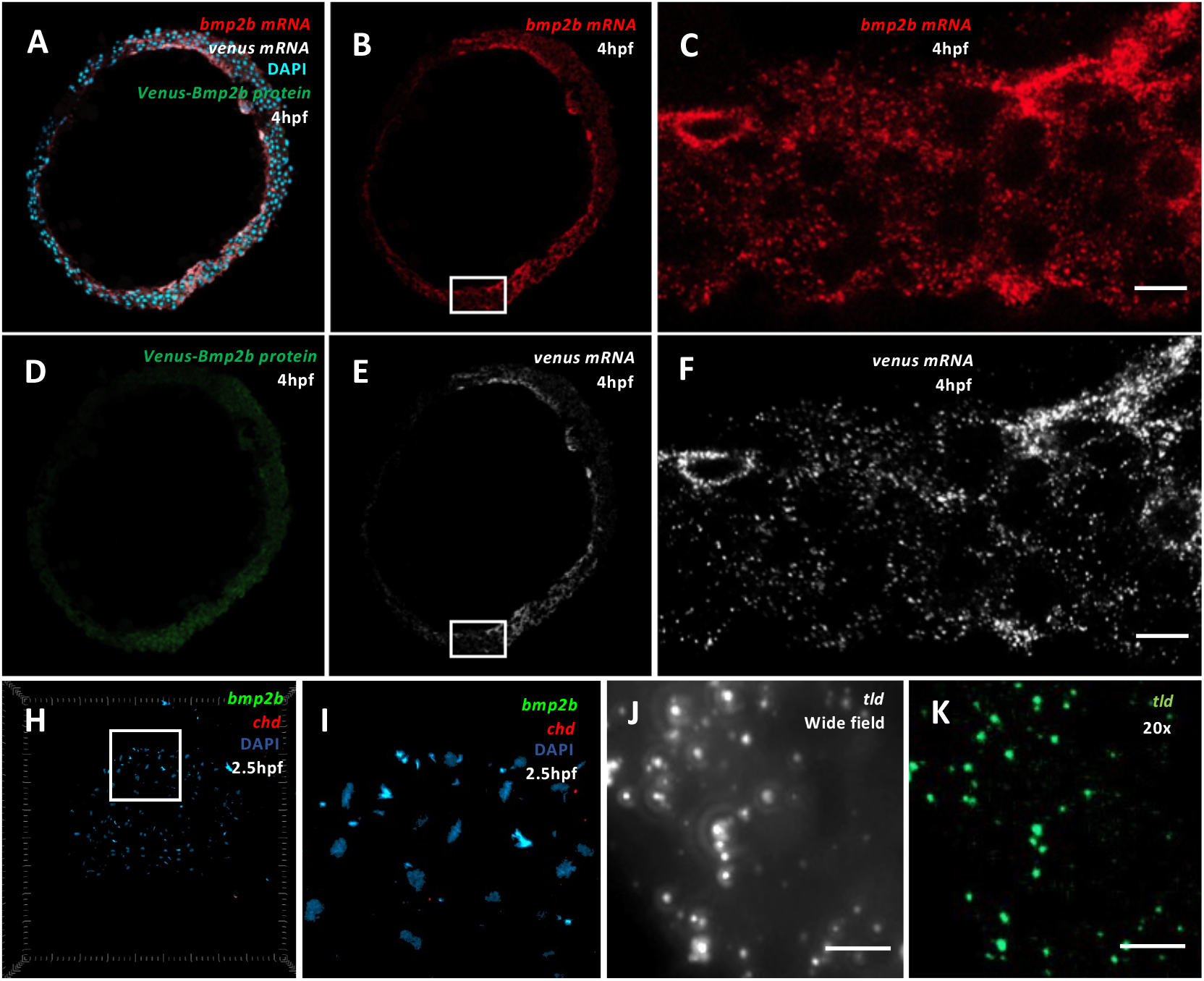
Specification of RNAscope probes in zebrafish. (A-F) (B-C) *bmp2b* mRNA(red), (E-F) *venus* mRNA(gray), DAPI (blue) and (D) Venus-Bmp2b protein(green) expression on zebrafish cryosection at 4hpf, into which *venus-bmp2b* fusion mRNA are injected at 1 cell stage. Scale bar: 10 μm. (C, F) The white box in B and E with higher magnification. (H-I) *bmp2b*(green), *chd*(red) and DAPI in zebrafish whole-mount embryos at 2.5hpf. (I) The white box in H with higher magnification. (J, K) *tld* mRNA imaged on wide-field microscope(J) and confocal 20x objective(K). Scale bar: 5μm

**Sup Fig.2.**
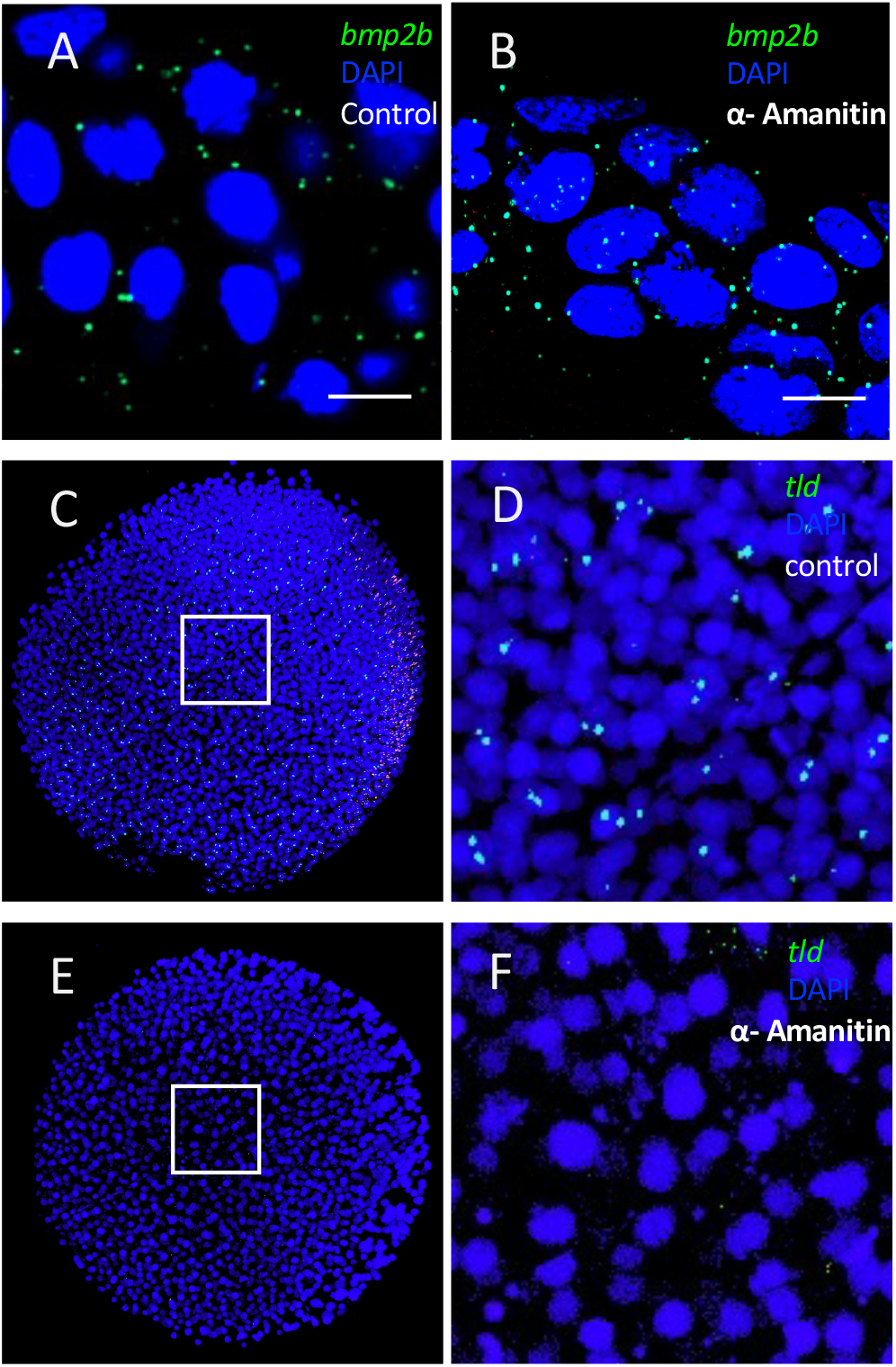
Transcription inhibition of *bmp2b* and *tld* mRNA in zebrafish at 6hpf. (A, B) *bmp2b* mRNA(green) in control(A) and embryos injected with alpha-amanitin(B).Scale bar: 10 μm. (C-F) *tld* mRNA(green) in control (C, D) and embryos injected with alpha-amanitin (E, F). (D, F) The white box in C and E at higher magnification.

**Sup Fig.3.**
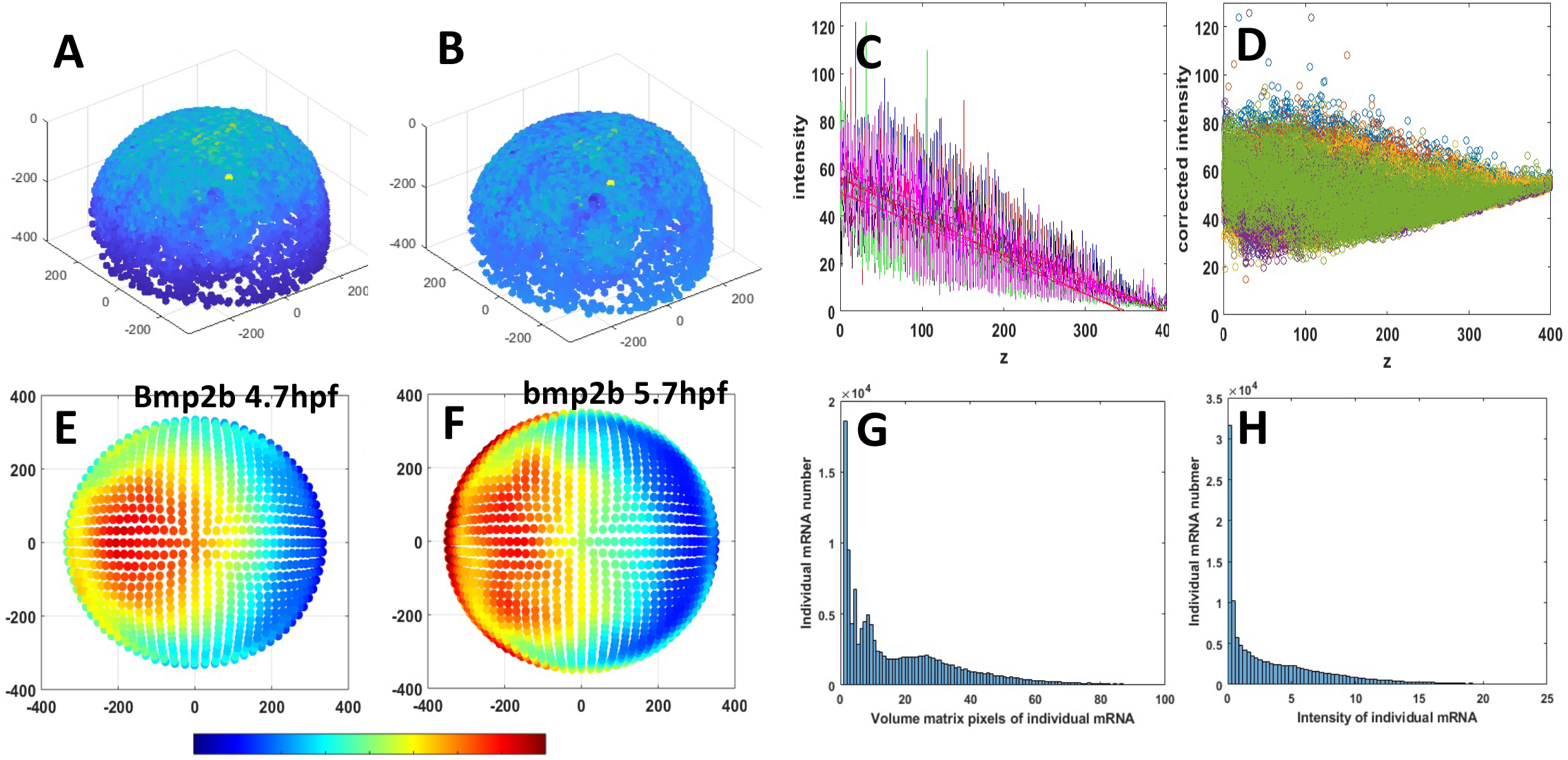
mRNA quantification related to Figure 2. (A-D) intensity drop-off correction in whole-mount embryos(n=5). (A) nucleus intensity at 647 channel decreased from animal pole to marginal region and quantification of averaged intensity at each slice are shown in (C).(B) nuclear intensity after correction and the corrected scatter points are shown in D.(E,F) Averaged *bmp2b* mRNA intensity distribution at 4.7hpf (E, n=3) and at 5.7hpf (F, n=4). (G-H) Statistical distribution of individual mRNA number at different volume matrix pixels(G) and intensity(H) of individual mRNA

**Sup Fig.4.**
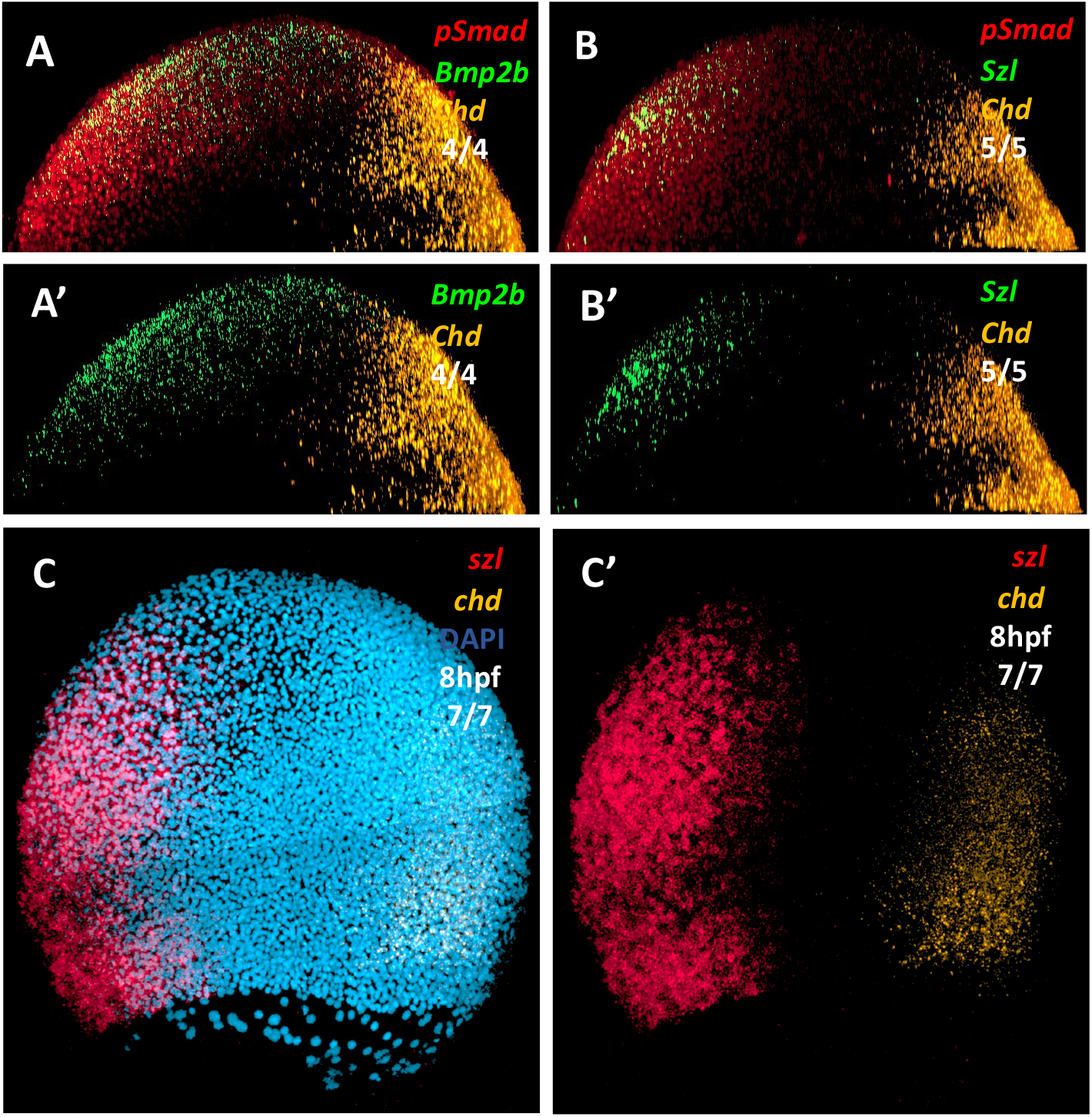
mRNA and protein expression from blastula to gastrula stage, related to Figure 3. (A-B’) simultaneous expression of pSmad (red), *bmp2b* (A, A’) or szl (B, B’) (green) and *chd* (orange) single molecular mRNA in lateral view at 5.7hpf. (C, C’) *szl*(red) and *chd* (orange) single molecular mRNA in lateral view at 8hpf with nucleus stained by DAPI (blue).

